# Dissociable, species-specific impact of Aβ on static and dynamic functional connectomes

**DOI:** 10.64898/2026.04.26.720907

**Authors:** Matteo M. Grudny, Nicholas Rodriguez, Thomas J. Murdy, Zachary D. Simon, Quan Vo, Wen Li, Matthew R. Burns, Damon G. Lamb, Catherine C. Kaczorowski, Paramita Chakrabarty, Marcelo Febo, the Alzheimer’s Disease Neuroimaging Initiative

## Abstract

Temporal dynamics in functional connectomes offer a physiologically grounded signature of ‘hidden’ pathologies during preclinical stages of Alzheimer’s disease (AD). We evaluated the effect of beta-amyloid (Aβ) on dynamic functional connectomes in transgenic mice and human subjects. Functional magnetic resonance images (fMRI) were collected in two strains of Aβ mice. fMRI-derived connectomes were segmented into discrete states using a hidden Markov model and network strength, efficiency, and transitivity were analyzed per state. Human fMRI-derived connectome measures were analyzed across 3 states. Static network measures were significantly different between Aβ mice and controls, the former having high values for strength, efficiency and clustering coefficient in anterior cingulate, hippocampus and retrosplenium. Dynamic network measures were stable within-states in Aβ mice. Similarly, human subjects with high Aβ had high node strength in precuneus and temporoparietal areas compared to low Aβ. In contrast, however, high Aβ was associated with high state switch rates, high fractional occupancy and state dwell times. Also, global strength, efficiency, and transitivity were less stable within states in the high Aβ group. Our results indicate that static, but not dynamic, connectome strength, efficiency and network integration are increased in Aβ mice, while dynamic network states appear less stable in human functional connectomes. This data supports a dissociable, species-specific impact of Aβ, with dynamic network alterations present in humans but not in Aβ mouse models, suggesting additional non-Aβ-driven influences on dynamic functional connectivity in preclinical AD.

**SIGNIFICANT STATEMENT:** Detecting Alzheimer’s disease before symptoms appear remains a critical challenge. This study reveals that beta-amyloid — a hallmark Alzheimer’s protein — disrupts brain network dynamics differently across species. While both transgenic mice and humans with elevated amyloid show stronger static brain connectivity, only humans exhibit unstable, rapidly shifting brain network states. This dissociation suggests that the dynamic network disruptions seen in preclinical Alzheimer’s patients are not solely driven by amyloid accumulation, but likely reflect additional biological factors absent in mouse models. These findings highlight an important translational gap between animal models and human disease, with implications for how we design and interpret preclinical Alzheimer’s research and develop early diagnostic biomarkers.

## INTRODUCTION

The expression of core Alzheimer’s disease (AD) symptoms is preceded by a protracted period in which known pathological hallmarks act along with ‘latent’ pathologies to progressively impair functional networks (Herrup, 2015; Masters et al., 2015). During this prodromal phase, the buildup of extracellular beta amyloid (Aβ) is central to functional deficits in neuronal circuits serving memory (Masters et al., 2015). Network-centric functional magnetic resonance imaging (fMRI) has helped uncover part of the brain regional interactions adversely affected by Aβ (Buckner et al., 2005; Buckner et al., 2009; Hedden et al., 2009; Wang et al., 2013; Zhang et al., 2024). Understanding how functional brain networks are reconfigured during preclinical AD could uncover novel pathological events to mark early-stage disease vulnerability(Pini et al., 2025).

Elevated Aβ is associated with altered functional connectivity in medial frontal, posterior cingulate, temporal-parietal and precuneus areas of the default mode network (DMN), and parahippocampal regions (Myers et al., 2014; Pereira et al., 2018; Fischer et al., 2025). Functional connectivity between precuneus and anterior hippocampus in individuals with high Aβ levels is increased, while a decline in functional connectivity is observed in normal aging individuals with low Aβ. Significant declines in functional connectivity across DMN areas are also reported in AD and mild cognitive impairment (MCI)(Wang et al., 2013; Dai and He, 2014; Dai et al., 2015), although this to a larger extent involves tau-mediated atrophic changes (Brier et al., 2014; Susanto et al., 2015; Canu et al., 2017; Jones et al., 2017; Cieri et al., 2022). Neuroimaging studies using transgenic rodents support a link between Aβ and increased functional connectivity involving frontal, cingulate, hippocampal and DMN-like regions(Shah et al., 2016; Latif-Hernandez et al., 2019), while other AβPP strains have age-progressive reductions in functional connectivity in parietal, motor, retrosplenial and dorsal hippocampal regions(Shah et al., 2013; Grandjean et al., 2014; Manno et al., 2019), particularly in the presence of tau (Manno et al., 2019).

The reconfiguration of functional interactions between network nodes may be key to AD progression (Cha et al., 2026). Amyloid positivity has been closely linked to node strength, clustering coefficient, and efficiency, although this has varied across several studies(Kesler et al., 2018; Munoz-Moreno et al., 2018; Simon et al., 2025). In addition, there is growing evidence that Aβ and other AD pathologies affect functional connectivity network measures across distinctly discernible temporal states(Rashid et al., 2014; Schumacher et al., 2019; Lin et al., 2020; Dautricourt et al., 2022; Qian et al., 2024). The temporal evolution of functional connectivity during a single session of imaging is thought to capture physiologically relevant dynamic states(Calhoun et al., 2014; Zalesky et al., 2014), perhaps internally driven by variability in cognitive and emotional neuronal circuit processing (Bassett et al., 2011; Nguyen et al., 2017). Dynamic networks have been studied in rodents and compared to human and non-human primates, with emerging features in common(Gutierrez-Barragan et al., 2024). Risk of AD and dementia, and progression to AD, are associated with temporal variability in functional connectivity (Dautricourt et al., 2022; Sendi et al., 2023a; Sendi et al., 2023b). Temporal variability in network connectivity is also observed in subjective cognitive decline, with pairwise regional changes in dynamic connectivity that are specific to cognitive decline (Wang et al., 2022).

In the present study, we used a variant of the hidden Markov model (HMM)(Vidaurre et al., 2017) to inference ‘hidden states’ underlying spatiotemporal functional connectome measures of integration and segregation in mice bearing Aβ, and in a group of ADNI data sets of elderly individuals with low versus high Aβ and no cognitive impairment. Both in mouse and in human data sets, Aβ is associated with high network strength, efficiency and clustering coefficient in DMN areas, and DMN-like areas in mice. However, while in mice we find that the effect of Aβ on these network measures persists stably within states, in humans we observe that high Aβ is linked to variable strength, efficiency and clustering coefficient within states. This cross-species difference suggests ‘hidden’ pathological mechanisms in human AD that are not present in mice only harboring Aβ pathology.

## MATERIALS AND METHODS

### Mice

This study used two strains of adult transgenic (TG) mice that develop amyloid plaques, TgCRND8 mice(Chishti et al., 2001) and a sub-strain of the recombinant inbred AD-BXD line(Neuner et al., 2019). Age, strain, and sex-matched non-transgenic (NTG) mice served as controls. Mice were housed in groups of 3-4 in a temperature- and humidity-controlled room, inside conventional air filtered cages (dimensions: 29 x 18 x 13 cm) with food and water available *ad-libitum* (vivarium lights on from 07:00-19:00 h).

Methods describing the development and maintenance of TG strains are published(Colon-Perez et al., 2019; Neuner et al., 2019; McFarland et al., 2021). TgCRND8 mice have early onset expression of human mutant beta-amyloid protein precursor (AβPP) (Swedish AβPP KM670/671NL and Indiana AβPP V717F), which increases human AβPP 5-times above endogenous murine AβPP. AD-BXD mice (referred to here as 5XFAD) are hemizygous for the dominant 5XFAD transgene(Oakley et al., 2006), which consists of 5 human mutations known to cause familial AD (three AβPP mutations: Swedish K670N, M671L, Florida I716V, and London V717I and two presenilin-1 PSEN1 mutations: M146L and L286V).

A total of 95 mice were used, including 7-8 month old (m) (young NTGY, n=19; 7 female mice), 15-19 m (middle age NTGM, n =9; 5 female mice), 21-23 m (aged NTGA, n = 25; 10 female mice) NTG mice, 6-9 m (young CRND8Y, n=13; 7 female mice) and 16-18 m (middle age CRND8M, n = 5; 3 female mice) CRND8 mice, and 7-8 m (young 5XFADY, n= 12; 5 female mice) and 15-18 m (middle age 5XFADM, n=12; 5 female mice) 5XFAD mice. TG mice were scanned at two timepoints, with TgCRND8 losing 8 mice by 16 m and 5XFAD mice showing 100% survival at 15 m. Littermates for the 5XFAD mice included NTG C57BL6/J (B6) and B6D2 (B6xDBA/2J F1). The background strain for TgCRND8 was B6C3H (B6xC3H/HeJ F1) obtained from the Jackson laboratory (Bar Harbor). Procedures were approved by the Institutional Animal Care and Use Committee of the University of Florida and follow all applicable NIH guidelines.

### Magnetic resonance imaging

Data were collected on an 11.1 Tesla MRI system equipped with Resonance Research Inc. gradients (RRI BFG-240/120-S6, 1000 mT/m at 325 Amps, 200 µs risetime) and a Bruker AV3 HD console running Paravision 6.0.1. A custom in-house quadrature transceive surface radiofrequency (RF) coil was used for image acquisition (Advanced Magnetic Resonance Imaging and Spectroscopy Facility, Gainesville, FL). Mice were imaged sedated under 0.1 mg/kg (i.p.) dexmedetomidine and a continuous paranasal flow of 0.25 % isoflurane (0.5 L/min flow; 70% nitrogen and 30% oxygen; Airgas, Inc.). An infusion line subcutaneously delivered supplemental dexmedetomidine during scanning (0.1 mg/kg/ml at an infusion rate of 25 µl/hour (PHD-Ultra microinfusing pump, Harvard Apparatus, Holliston, MA). Functional MRI scans were collected at least 50 minutes after the dexmedetomidine injection. Mice were kept warm during scanning and spontaneous breathing rate was monitored.

A T2-weighted Rapid Acquisition with Refocused Echoes (RARE) sequence was acquired with the following parameters: echo time (TE) = 41 ms, repetition time (TR) = 4 seconds, RARE factor = 16, number of averages = 12, field of view (FOV) of 15 mm x 15 mm and 0.9 mm thick slice, and a data matrix of 256 x 256 (0.06 mm^2^ in plane) and 14 interleaved ascending coronal (axial) slices, covering the entire brain from the rostral-most extent of the anterior prefrontal cortical surface, caudally towards the upper brainstem and cerebellum. Functional images were collected using a single-shot spin echo planar imaging (EPI) sequence with the following parameters: TE = 15 ms, TR = 2 seconds, 600 repetitions, FOV = 15 x 15 mm and 0.9 mm thick slice, and a data matrix of 64 x 48 (0.23 x 0.31 mm in plane) with 14 interleaved ascending coronal slices in the same position as the anatomical scan. Ten dummy EPI scans were run prior to acquiring data. Respiratory rates, isoflurane and dexmedetomidine delivery, temperature, lighting, and room conditions were kept constant across subjects.

### Image processing

Images were processed using Analysis of Functional NeuroImages (AFNI)(Cox, 1996), FMRIB Software Library (FSL)(Jenkinson et al., 2012), and custom scripts written in MATLAB(Febo et al., 2024). Voxel time series were ‘despiked’, motion-corrected, and drift-corrected. We ran independent components analysis (ICA)(Beckmann and Smith, 2004) to evaluate time series and generate nuisance regressors. These nuisance signals were identified by the presence of scattered or individual voxels with large irregular and spurious temporal shifts, high frequency and/or pulsatile temporal patterns, mostly localized to clusters of voxels along brain edges and spurious voxels in white matter areas and ventricles. A list of nuisance regressors was generated per mouse. fMRI scans were ‘denoised’ using these regressors, band pass filtered to suppress frequencies outside of a bandwidth of 0.009 Hz to 0.2 Hz and spatially filtered. This frequency bandwidth allowed a broad range of signal features to explore dynamic network states (see below). A functional image volume was used for multi-subject fMRI template-construction using Advanced Normalization Tools (ANTs)(Klein et al., 2009). The single fMRI volume per subject was bias field corrected, and brain extracted before template construction. Affine and nonlinear transformations were applied to preprocessed fMRI scans, and these were analyzed in template space. Inverse linear transform of a mouse atlas(Johnson et al., 2010) parcellation to the multisubject template was performed to create 51 regions of interest (ROI) masks per hemisphere (102 total bilateral ROI masks) to extract time series.

### Static networks

Pearson correlations between pairs of ROIs (totaling ∼102^2^) were carried out in MATLAB. The r coefficients were Fisher z transformed, organized into weighted undirected symmetrized matrices, and graph calculations for a network density of 15% carried out using Brain Connectivity toolbox(Rubinov and Sporns, 2010). These included network strength, characteristic path length, global efficiency, assortativity, modularity and transitivity(Febo et al., 2024). *Node strength* was calculated as the sum of edges (Pearson r’s) per ROI, with the global average across ROIs used as an indicator of network strength. *Characteristic path length* was calculated by first converting correlation matrices to length matrices and then using Dijkstra’s algorithm to estimate distances. Thus, characteristic path length was the mean weighted distances across all nodes and *global efficiency* was the inverse distance matrix. For *transitivity*, correlation matrices were scaled to 0,1 and the ratio of triadic groups (strongly interconnected groups of 3 nodes) normalized to all possible triplets was used as a measure of how well integrated nodes are within the network. Node level versions for transitivity (clustering coefficient) and global efficiency (node efficiency) were also calculated. *Assortativity* was calculated as the correlation between strength values of above-diagonal row and column elements of adjacency matrices. *Modularity* was calculated using the classical optimal community structure algorithm that indexes the degree of maximum within-group and minimal between-group edges. This was used as an index of segregation of nodes into functional communities. Matrices were averaged within experiment groups, mean matrices set to an edge density of 15%, node strengths calculated and then visualized in BrainNet Viewer with nodal spheres overlaid onto a mouse atlas 3-dimenstional translucent shell representing node strength and the interconnecting lines the above-threshold edge weights, as previously described(Febo et al., 2024).

### Dynamic networks

Formal description of HMMs applied to fMRI time series has been described (Vidaurre et al., 2017). Briefly, an HMM segments multivariate time series into a sequence of discrete “states” with Markov property. For a consecutive series of observations, X_1:T_, and hidden states, Z_1:T_ ∈ {1,…,K}, with joint distribution 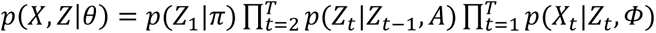 (Bishop, 2006), the model parameters are denoted by the term θ = {π, A, Φ}, where π is the initial state distribution, A is the K×K transition matrix with row sums equal to 1, and Φ are the emission parameters(Bishop, 2006). Here we used Gaussian emissions with MAR order 0 (no autoregression; thus, Φ = one mean and covariance per state) after standardization (z-scoring) and dimensionality reduction (PCA) of windowed connectivity features. Inference is performed using variational Bayes, as implemented in HMM-MAR, which maximizes the variational free energy (evidence lower bound)(Bishop, 2006). A Dirichlet prior (controlled by the DirichletDiag parameter in HMM-MAR, which we set to 100(Vidaurre et al., 2017)) is placed on each row of A for reasonable self-transitions. This yields posterior state probabilities per time point, Gamma, Γ_t,k_ = p(Z_t_ = K | X, θ), and the most probable state sequence via the Viterbi algorithm. Representative ‘state mean networks’ for each HMM state was estimated by Γ-weighted averaging of connectivity windows that were mapped back to ROI×ROI matrices. For each state K, we computed a mean edge vector, 𝑀_𝐾_ = (∑_𝑡_ 𝛤_𝑡,𝐾_ · 𝑋_𝑡_) / (∑_𝑡_ 𝛤_𝑡,𝐾_), where X_t_ is the vectorized upper-triangle of the windowed Fisher-z connectivity and Γ_t,K_ is the posterior probability of state K at time t. We then filled M_K_ into the upper triangle, symmetrized, and set the diagonal to zero to obtain the state’s mean adjacency, G^K^. We derived dynamic features (fractional occupancy, mean dwell time, and the empirical transition matrix) from Γ and the corresponding most probable state (MAP/Viterbi) sequence. Graph-theoretic measures per state were computed on the state mean adjacencies, G^K^, after proportional thresholding to a 15% graph density.

The schematic in **Figure 1** summarizes the preprocessing steps in HMM-MAR. We imported ROI time series into MATLAB (600 time points by 102 ROIs) and applied a sliding window (30 TR, 1 TR step, so 60-second segments with 2 second stride) to generate 571 windows ([timepoints – windows] / step size + 1). For each window we computed pairwise correlations, Fisher z-transformed them, and vectorized the upper triangle ([102 x 101]/2] for a total of 5,151 edges. The result is a 571 x 5,151 window-by-edge matrix per subject (later concatenated across subjects) as input to HMM-MAR. We fit a Gaussian HMM with shared full covariance across states, standardization, and PCA (retaining ∼52% explained variance). We used 15 restarts (initrep) with 25 initialization cycles (initcyc) and allowed up to 1,000 inference cycles. We swept K = 3-10 (in line with prior rodent work(Tsurugizawa and Yoshimaru, 2021; Gutierrez-Barragan et al., 2022; Gutierrez-Barragan et al., 2024), although we note other reports inference a greater number of states(Bukhari et al., 2017; Grandjean et al., 2017)) and selected K=5. At higher K (6–10) increasingly many ‘microstates’ with negligible durations (<1–2 TR) were observed.

**Figure 1.**
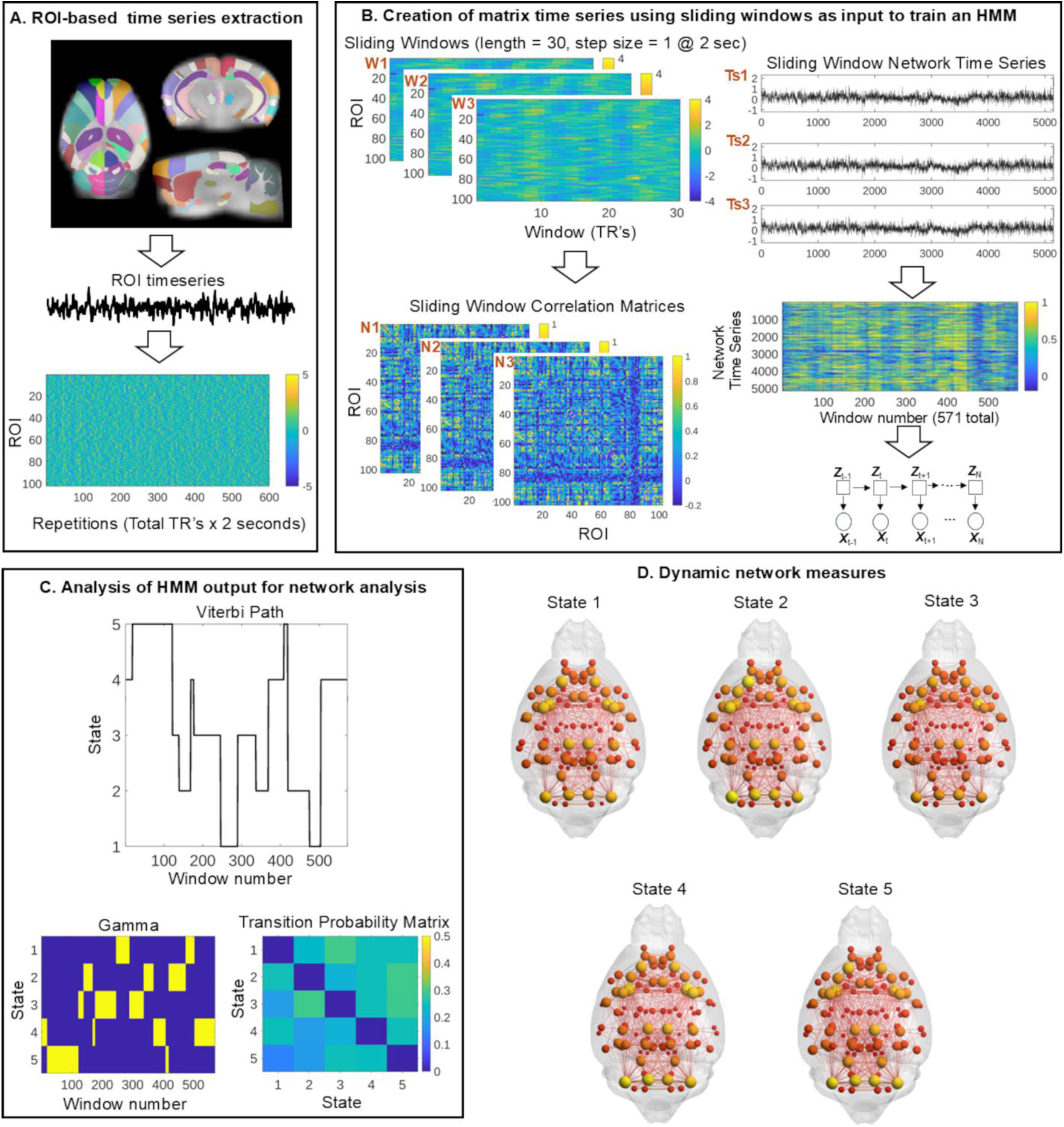
Processing of mouse fMRI signals for HMM-MAR. A) Signals were extracted from post-processed images aligned to a mouse brain parcellation(Johnson et al., 2010). B) Following a sliding window, network matrices were constructed per window and vectorized and organized in a network x window matrix as input to HMM-MAR(Vidaurre et al., 2024). C) HMM outputs. D) Network visualizations across states.

### Histology

Aβ plaques were quantified in hemi-brains of TgCRND8 mice (n=5; 6m mice n = 3, 12m mice n = 2 mice). Brain harvesting was done at the University of Florida and tissue preservation, processing, clearing, immunolabeling and volumetric imaging of immunolabeling, and image post-processing was carried out at Lifecanvas Technologies (Cambridge, MA). Paraformaldehyde-fixed brains were preserved using SHIELD reagents(Park et al., 2018), delipidated and labeled using eFLASH(Yun et al., 2025) technology on a SmartBatch+ device. Aβ plaques were co-labeled with propidium iodide (PI) nuclear stain as background. Immunolabeling was carried out with Aβ anti-body (mouse IgG2a, MCA-AB9, Encor) and donkey anti-mouse Alex Fluor 488 secondary. After immunolabeling, samples were incubated in EasyIndex (RI = 1.52) for refractive index matching. Intact whole brain volumes were imaged using a SmartSPIM axially swept light sheet microscope using a 3.6x field of view (0.2 NA objective) at 20 fps during volumetric acquisition with 488 and 561 nm laser lines for Aβ and PI. Registration to the mouse common coordinate framework v3 (Allen Mouse Brain Atlas) followed published pipelines(Perens et al., 2021), SmartAnalytics code was used for Aβ quantification and image processing (Perens et al., 2021) and analysis of percent Aβ occupation per ROI(Goubran et al., 2019) (LifeCanvas Technologies, Cambridge, MA).

### Static and dynamic functional networks in low and high amyloid density

Data used in the preparation of this article were obtained from the Alzheimer’s Disease Neuroimaging Initiative (ADNI) database (adni.loni.usc.edu). The ADNI was launched in 2003 as a public-private partnership, led by Principal Investigator Michael W. Weiner, MD. The primary goal of ADNI has been to test whether serial magnetic resonance imaging (MRI), positron emission tomography (PET), other biological markers, and clinical and neuropsychological assessment can be combined to measure the progression of mild cognitive impairment (MCI) and early Alzheimer’s disease (AD).

We analyzed static and dynamic functional networks in an Alzheimer’s disease neuroimaging initiative (ADNI3) dataset (approved under University of Florida protocol #: NH00047739). Out of 34 scans, 27 were controls and 7 had subjective memory complaints. None were diagnosed with mild cognitive impairment or Alzheimer’s. The dataset included 16 subjects with high amyloid density (age 72.2 ± 7.0; 2 males; 1.41 ± 0.19 AV45 SUVR and 1.38 ± 0.17 at baseline) and 18 with low amyloid density, defined by AV45 normalized binding (age 69.2 ± 6.2; 2 males; 1.00 ± 0.03 AV45 SUVR and 1.01 ± 0.04 at baseline)(t-test: low vs high Aβ, *p* = 4.4 x 10^-7^). Low and high Aβ groups all had CDRSB scores of 0 with no evidence of cognitive impairment from their Montreal Cognitive Assessment scores (MoCA: low Aβ = 27.2 ± 1.69, high Aβ = 25.6 ± 2.9; t-test *p* = 0.011), mini-mental state exam scores (MMSE: low Aβ = 29.38 ± 0.69, high Aβ = 28.5 ± 1.55; t-test *p* = 0.04), and Alzheimer’s dementia assessment scale scores (ADAS-Cog11: low Aβ = 5.79 ± 2.1, high Aβ = 6.4 ± 4.2; t-test *p* = 0.6; ADAS-Cog13: low Aβ = 8.62 ± 3.2, high Aβ = 9.85 ± 6.1; t-test *p* = 0.48).

Axial EPI scans were collected on 3 Tesla Siemens scanners (70% Prisma, 21% Verio, and 9% Skyra) with the following parameters: TE=30ms, TR=3seconds, dimensions 3.4 mm^3^, matrix size = 64x64x48 voxels, 197 repetitions (10-minute scan duration). Subjects had eyes open during fMRI scanning. The first 7 volumes in the 197 volume time series were removed prior to processing steps. The steps included time series spike removal, slice timing correction, motion and drift correction. ICA based identification of ‘noisy’ time series voxels was used to regress these prior to temporal filtering between 0.009 and 0.2 Hz, and spatial filtering. A temporal average volume was used to create a multi-subject template guided by affine and non-linear normalization to the Montreal Neurological Institute template. The Schaeffer 300 parcellation(Schaefer et al., 2018) was used to extract ROI time series. Adjacency matrices comprising Fisher-z transformed pairwise correlations ([300 * 299) / 2] = 44,850 edges) were set to 20% density prior to network calculations. These included the same measures described above. Weighted undirected networks were visualized in BrainNetViewer (Xia et al., 2013).

Network time series were generated and used in HMM-MAR inference as described above and as illustrated in **Figure 2**. ROI time series were imported into MATLAB (190 time points by 300 ROIs) and a sliding window applied (30 TR, 1 TR step, for 90-second segments at 3 strides) to generate [timepoints – windows / step size + 1] 161 connectivity windows. Matrices were Fisher transformed and reorganized to 44,850 edges by 161 windows for HMM-MAR. HMM-MAR parameters were as described above. We conducted iterative inferencing of K from 3-7, selecting 3 based on the same criteria as described above. Dynamic features and graph theory network estimates for a 20% density threshold were derived as described above. As with mouse dynamic connectivity, we analyzed FO and dwell windows.

**Figure 2.**
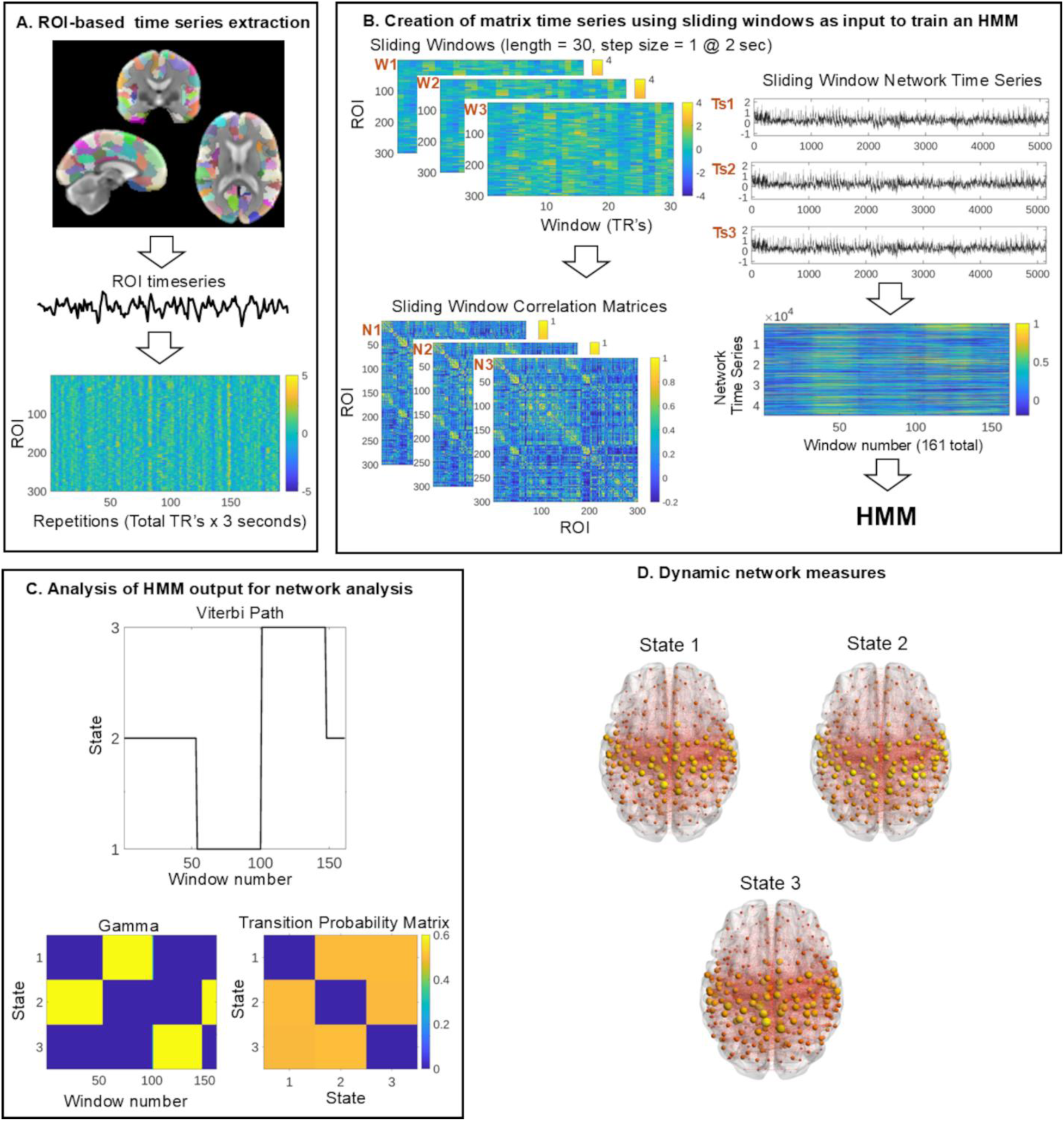
Processing of ADNI data-derived fMRI signals for HMM-MAR. A) Signals were extracted from post-processed images aligned to the Shaeffer 300 parcellation(Johnson et al., 2010). B) Following a sliding window, network matrices were constructed per window and vectorized and organized in a network x window matrix as input to HMM-MAR(Vidaurre et al., 2024). C) HMM outputs. D) Network visualizations across states.

### Statistical analysis

Statistical analyses were conducted in MATLAB (MathWorks, Inc, Natick, MA) and GraphPad Prism (Boston, MA). We assumed heteroscedasticity, with all groups in both mouse and human portions of the work having an unequal number of subjects. In the specific case of the mouse imaging studies, we had unbalanced groups, with control mice having 3 age conditions (young, middle aged and aged) and the TG strains having only young and middle-aged conditions.

For static networks, we analyzed global metrics using a non-parametric Kruskal-Wallis analysis of variance (ANOVA) with group-wise post hoc tests using Dunn-Sidak’s correction. For node level analysis, we used permutation-based statistical comparisons as implemented in Permutation Analysis of Linear Models (PALM)(Winkler et al., 2014). In addition to reasonable handling of multiple nonparametric contrasts, PALM handles both corrections for multiple comparisons across ROI’s as well as multiple group-level contrast corrections. A linear model was constructed with the FSL GLM tool to inference age, Aβ and their interaction. Specifically, we tested: (1) main effect of age by contrasting aged vs middle-aged and young mice regardless of strain, (2) main effect of Aβ through contrasts between Aβ mice vs NTG mice of any age, and 3 additional contrasts designed to test age x Aβ interactions independently for 5XFAD and CRND8 mice. We hypothesized age-related reductions in connectivity and amyloid related increases in connectivity across sensorimotor and cognitive areas. Permutations were done over the entire dataset, with the following options: Fisher non-parametric contrasts with family-wise error correction for number of contrasts, two-tail distributions, data demeaning, accelerated tail estimation, with 10,000 total shufflings, and all resulting log-P values were false discovery rate (FDR) adjusted across ROIs. Contrast T statistics are reported here along with FDR corrected p values. For rejections of the null hypothesis per ROI, post hoc tests between group ROI values were then carried out using Dunn-Sidak tests.

Dynamic functional connectivity state features and graph measures per state were analyzed with linear mixed effects (LME) ANOVA (MATLAB *lme*). For mouse dynamic networks, we tested for the main effect of group (7 groups: NTG, TG, subdivided by age), main effect of state (5 states), and group x state interaction (either global or nodal level network measure ∼ group*state + 1|subjects, where network measure is either global or nodal strength, efficiency or clustering coefficient). All resulting p-values for node-level comparisons were FDR adjusted to q ≤ 0.05 per term (main effect or interaction p values).

Human static and dynamic network comparisons, either at the global or nodal levels, followed the same statistical steps as the mouse networks, except for the following: (1) Mann-Whitney tests were used to compare low vs. high Aβ groups, (2) LME ANOVA compared 2 groups (either global or nodal level network measure ∼ group*state + 1|subjects, where network measure is either global or nodal strength, efficiency or clustering coefficient), (3) node level comparisons were done for nodes within DMN areas, according to the Schaeffer 300 node parcellation for 17 networks(Schaefer et al., 2018).

## Results

### Amyloidosis alters functional network strength, transitivity, and global efficiency

We first analyzed the effect of Aβ on static network measures in young and middle aged TgCRND8(Chishti et al., 2001) and 5XFAD(Neuner et al., 2019) transgenic (TGY, TGM) mice. These were compared to background young, middle aged, and aged non-transgenic (NTGY, NTGM, NTGA) controls. The network measures included network strength, characteristic path length, global efficiency, assortativity, modularity and transitivity(Febo et al., 2024). These graph measures were chosen to evaluate global functional network integration, segregation, and efficiency, that are affected in Alzheimer’s disease(Dai and He, 2014) and in amyloid mouse models(Kesler et al., 2018).

Network strength, clustering coefficient, and global efficiency significantly differed between the groups (Kruskal-Wallis ANOVA, p<0.05; **Figure 3**). Post hoc Dunn-Sidak tests did not reveal significant pairwise differences, although non-significant trends towards increased strength, clustering and efficiency were observed in TG strains compared to NTG (**Figure 3**).

**Figure 3.**
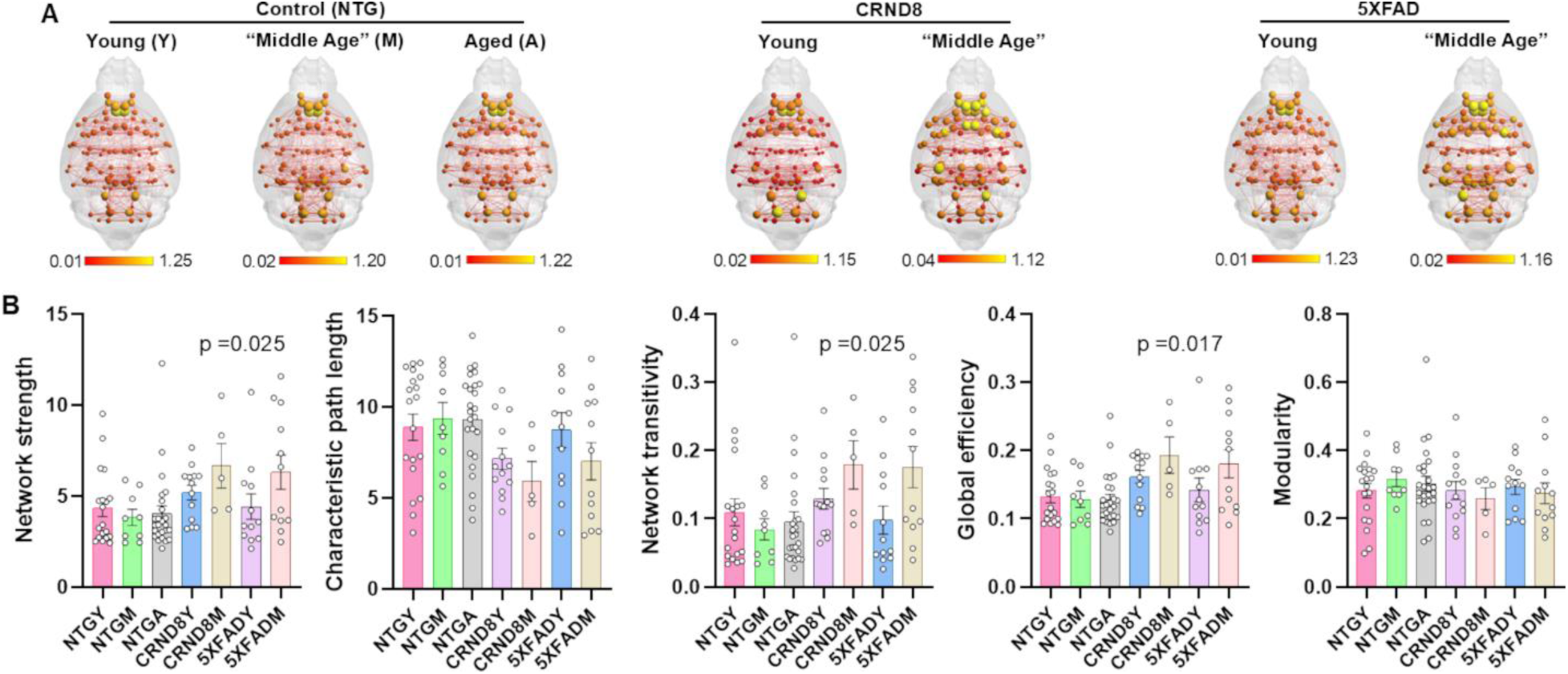
Amyloidosis, not age, alters global network strength, efficiency, and transitivity. A) Connectome maps of young (Y), middle age (M), and aged (A) non-transgenic (NTG) and transgenic (TG: TgCRND8, 5XFAD) mice. Maps visualized on mouse brain template(Johnson et al., 2010) with node intensity indicating strength of connectivity and lines (edges) indicating Pearson correlation (edge weights). Scale bar indicates rescaled (0,1) edge weights. B) Global network measures. Data presented as mean ± standard error with overlaid scatter plots. Significant group differences, p<0.05 (Kruskal-Wallis test).

Nonparametric contrasts (FDR-adjusted) comparing node strength between young, middle aged and aged NTG and young and middle-aged TG groups showed a significant main effect of Aβ in anterior cingulate cortex (t_89_ = 17.8, p=0.045), agranular insular cortex (t_89_ = 17.7, p = 0.045), secondary motor cortex (t89 = 15.6, p = 0.049), superior colliculus-motor area (t_89_ = 17.9, p = 0.045) in the left hemisphere, and superior colliculus-motor area (t_89_ = 16.1, p = 0.049), inferior colliculus (t_89_ = 19.9, p = 0.042), floccular nodal region of the cerebellum (t_89_ = 16.0, p = 0.049) in the right hemisphere (**Figure 4**). An age x Aβ interaction was observed in the retrosplenial cortex (t_89_ = 15.9, p = 0.033), primary visual area (t_89_ = 18.6, p = 0.031), subicular region (t_89_ = 17.2, p = 0.031), and periaqueductal gray (t_89_ = 18.4, p = 0.031) in the right hemisphere. Main effect of age was observed in the right periaqueductal gray (t_89_ = 16.9, p = 0.045). No age x amyloid interactions for each Aβ strain independently were observed. Post hoc (Dunn-Sidak) pairwise contrasts are summarized in **Figure 4A** and **Supplemental Figure 1A**.

**Figure 4.**
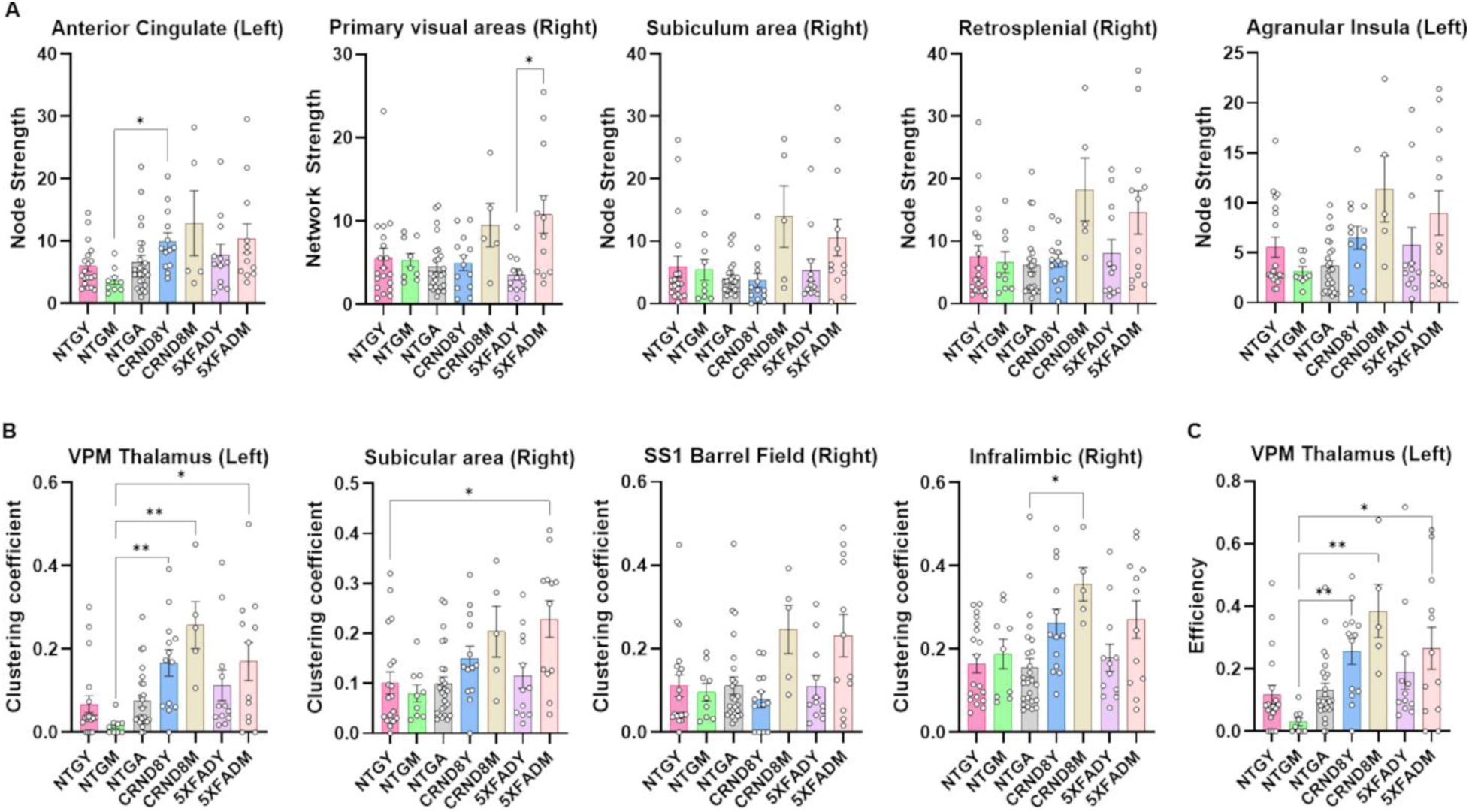
Amyloidosis alters node strength, clustering coefficient and efficiency across cognition and emotion related brain regions. Groups are as shown in Figure 2. A) Node strength. B) Clustering coefficient. C) Efficiency. All data presented as mean ± standard error with overlaid scatter plots. Significant differences tested across all nodes using linear mixed effects ANOVA (FDR corrected). Post hoc Dunn’s tests indicated by asterisks (*p<0.05, **p<0.01).

Nonparametric contrasts (FDR-adjusted) comparing clustering coefficients between groups showed a significant effect of Aβ in ventroposteromedial (VPM) (t_89_ = 29.9, p = 0.00069) and spinal nucleus of the trigeminal region (SpVr) (t_89_ = 19.6, p = 0.026) in left hemisphere and infralimbic cortex (t_89_ = 17.6, p=0.029) and subicular region (t_89_ = 19.6, p=0.029) in the right hemisphere. A significant age x Aβ interaction was observed in primary somatosensory area (t_89_ = 16.1, p = 0.046), and motor-related superior colliculus region (t_89_ = 16.1, p = 0.046) in the right hemisphere. No effect of age or age x amyloid interactions for each Aβ strain independently were observed. Post hoc (Dunn-Sidak) pairwise contrasts are summarized in **Figure 4B** and **Supplemental Figure 1B**.

Nonparametric contrasts (FDR-adjusted) comparing node efficiency between groups showed a significant effect of Aβ in VPM thalamus (t_89_ = 29.9, p = 0.001) in the left hemisphere. No significant age effect nor age related effect of Aβ was observed. No effect of age or age x amyloid interactions for grouped Aβ mice or for each Aβ strain independently were observed. Post hoc (Dunn-Sidak) pairwise contrasts are summarized in **Figure 4C**.

### Age and amyloidosis are associated with fluctuations in network strength, transitivity, and efficiency across functional connectome states

We used the MATLAB version of the hidden Markov model multivariate autoregressive (HMM-MAR) toolbox, published by Vidaurre(Vidaurre, 2021), to segment fMRI time series into quasi-stationary network states(Vidaurre et al., 2017). Using this approach, we evaluated the effect of age and Aβ on dynamic functional connectivity networks in mice. From these dynamic states, we analyzed fractional occupancy (FO: fraction of total time spent in each state) and dwell windows (number of windows per each state) as measures of dynamic functional connectivity variability. The average fractional occupancy (FO) per state across all groups was 0.20 ± 0.008 and the mean number of dwell windows per state was 41.8 ± 0.93.

LME ANOVA revealed a significant state and state x group effect on state transition probabilities (state effect F_19,1900_ = 1.75, p = 0.02; state x group interaction F_114,1900_ = 1.35, p = 0.008). No differences in FO and dwell windows were observed (**Figure 5**). We did not observe differences in state switch rates.

**Figure 5.**
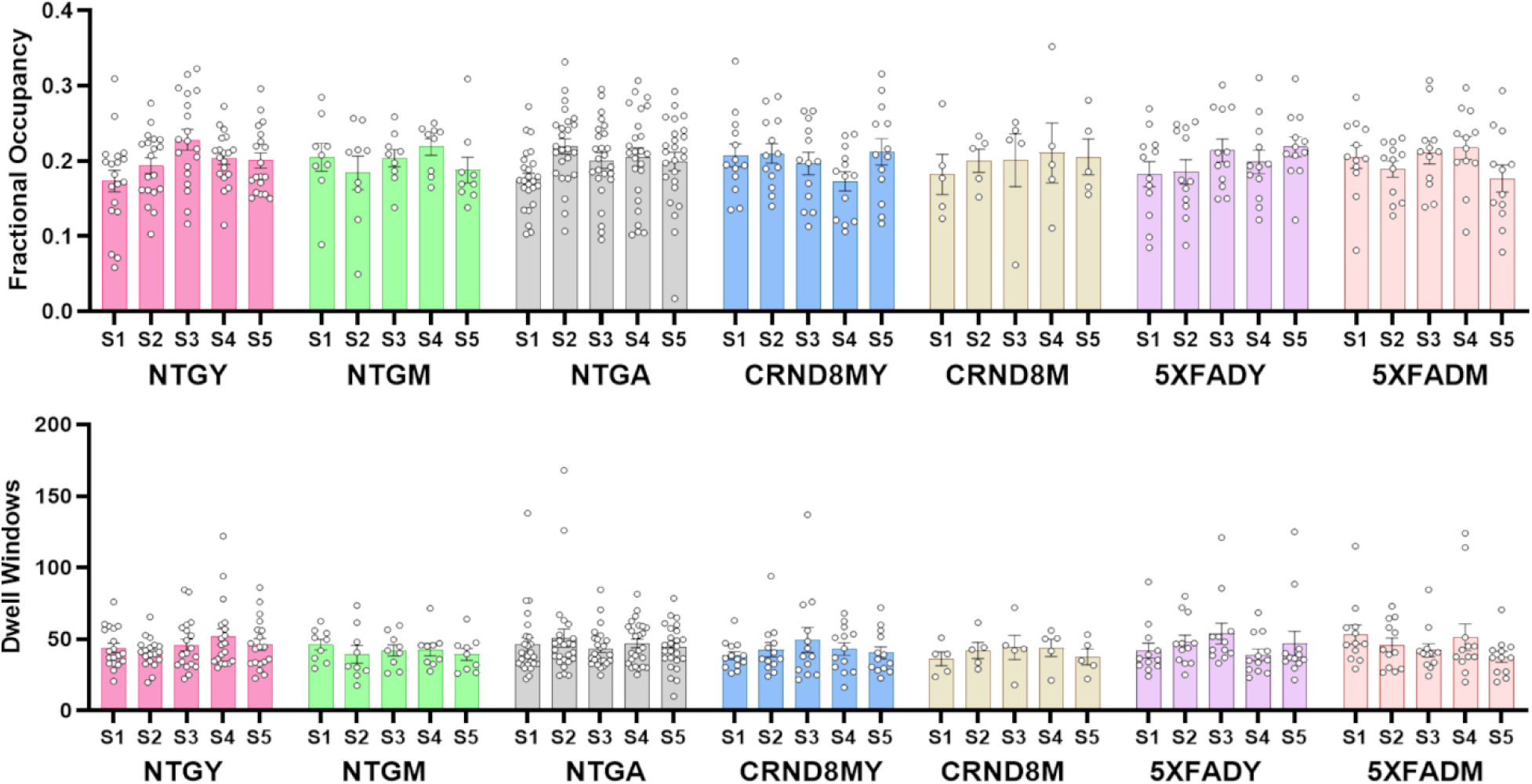
Stable dynamic functional network states in NTG and TG mice. Shown are fractional occupancy (fraction of total time spent in each state) and dwell windows (number of windows per each state). Groups are as shown in Figure 2. All data presented as mean ± standard error with overlaid scatter plots. Significant differences tested across all nodes using mixed effects ANOVA (FDR corrected). Post hoc Dunn’s tests indicated by asterisks (*p<0.05).

A 2-way mixed effect (ME) ANOVA indicated a main effect of group (F_6, 88_ = 2.6, p = 0.025) and group x state interactions (F_22.5, 329.8_ = 2.6, p = 0.0001) on network strength. Pairwise comparisons using Tukey’s multiple comparison test show within group (state fluctuations, specific to Aβ groups) and between groups differences in network strength. These are summarized in **Figure 6**. Increased network strength in Aβ groups, observed in static networks, persisted across dynamic networks.

**Figure 6.**
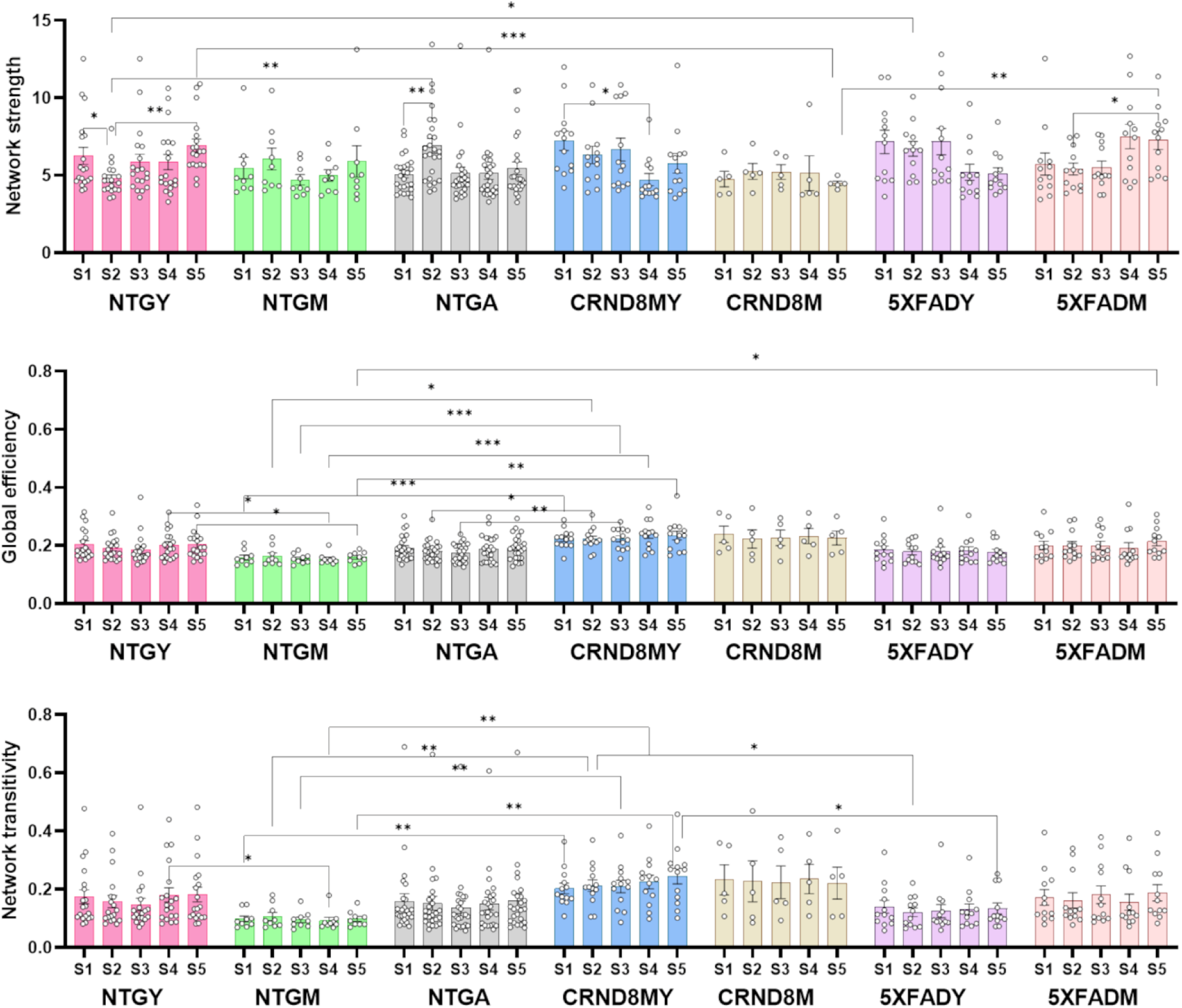
Amyloid associated group differences in strength, transitivity and global efficiency observed in static networks persist in dynamic functional networks. Groups are as shown in Figure 2. All data presented as mean ± standard error with overlaid scatter plots. Significant differences tested across all nodes using mixed effects ANOVA (FDR corrected). Post hoc Dunn’s tests indicated by asterisks (*p<0.05, **p<0.01, ***p<0.005).

We observed main effects of state (F_3.7, 321.5_ = 2.5, p = 0.049) and group (F_6, 88_ = 3.9, p = 0.002) on global efficiency (2-way ME ANOVA; **Figure 6**). No significant group x state interactions were observed. Pairwise comparisons using Tukey’s multiple comparison test show within group (state fluctuations, specific to Aβ groups) and between groups differences in global efficiency. These are summarized in **Figure 6**.

We also observed main effects of state (F_3.7, 321.5_ = 2.5, p = 0.048) and group (F_6, 88_ = 2.3, p = 0.04) on network transitivity (2-way ME ANOVA; Figure 5). No significant group x state interactions were observed. Pairwise comparisons using Tukey’s multiple comparison test show within group (state fluctuations, specific to Aβ groups) and between groups differences in transitivity. These are summarized in **Figure 6**.

LME ANOVA indicated a significant effect of state and a group x state interaction on node strength across several ROI nodes. The nodes are summarized in **Figure 7A**. The nodes with group x state interactions included left anterior cingulate (F_24,380_=3.3, FDR p = 0.00007), left anterior olfactory nucleus (AON) (F_24,380_=2.2, FDR p = 0.02), left dorsal striatum (F_24,380_=2.8, FDR p = 0.0008), left insula (F_24,380_=2.3, FDR p = 0.02), left supplemental somatosensory area (F_24,380_=2.1, FDR p = 0.02), left red nucleus (F_24,380_=2.2, FDR p = 0.02), and right basomedial amygdala (F_24,380_=2.6, FDR p = 0.003). The nodes with main effect of state included the left substantia innominata (F_4,380_=6.2, FDR p = 0.002), left central amygdala (F_4,380_=6.6, FDR p = 0.001), left auditory cortex (F_4,380_=8.3, FDR p = 0.001), left PAG (F_4,380_=3.9, FDR p = 0.00007), right superior colliculus (F_4,380_=4.5, FDR p = 0.02), left (F_4,380_=4.7, FDR p = 0.02) and right basomedial amygdala (F_4,380_=9.1, FDR p = 0.00005) and left (F_4,380_=3.9, FDR p = 0.04) and right cerebellar vermis (F_4,380_=4.8, FDR p = 0.02).

**Figure 7.**
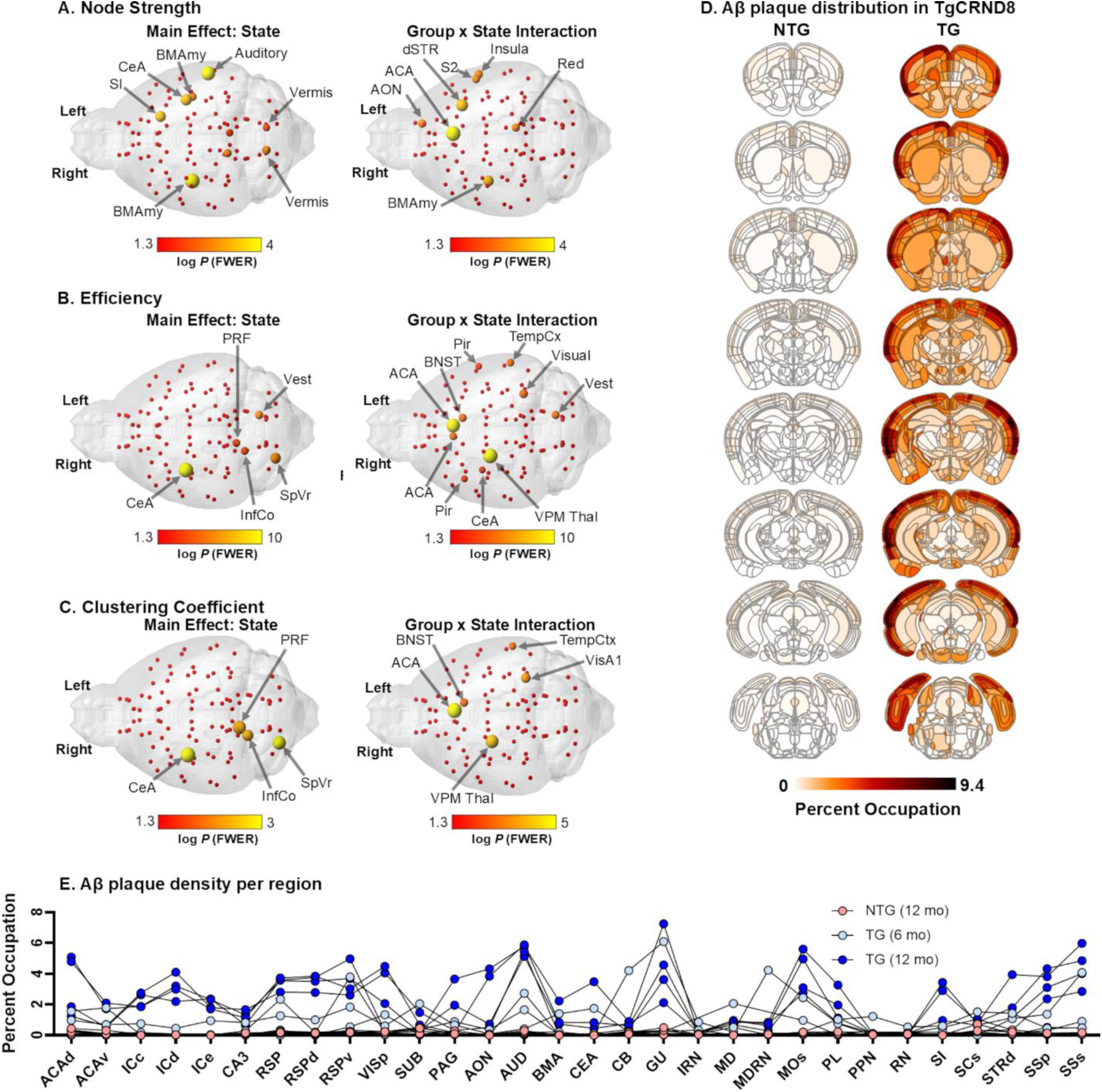
Group differences in node strength, efficiency and clustering coefficient vary across dynamic states in cognition and emotion related brain regions. A-C) Nodes showing main effect of state and group x state interactions for node strength (A), efficiency (B), and clustering coefficient (C). Scale bar color intensity indicates log p value. Significant differences tested across all nodes using linear mixed effects ANOVA (FDR corrected). D) Whole brain histology of Aβ plaque distribution in brains of an NTG and a TgCRND8 mouse. Scale bar indicates percent occupation of Aβ signal per ROI. E) Aβ plaque density per region. Data presented as individual data points per hemibrain (NTG, n=2; 6mo TG, n=3, 12mo TG, n=2). Abbreviations: ACA, anterior cingulate; IC, internal capsule; CA3, hippocampus; RSP, retrosplenium; VIS, visual cortex; PAG, periaqueductal grey; AON, anterior olfactory nucleus; AUD, auditory cortex; BMA, basomedial amygdala; BNST, bed nucleus stria terminalis; CEA, central amygdala; CB, cerebellum; GU, gustatory cortex; IRN, intermediate reticular nucleus; MD, mediodorsal thalamus; MDRN, midbrain reticular nucleus; MO, motor cortex; Pir, piriform; PL, prelimbic, RN, reticular nucleus; PPN, pedunculopontine nucleus; SI, substantia innominata; SC, superior colliculus; STR, striatum, SS, somatosensory; SpVr, spinal trigeminal nucleus region; VPM thal, ventroposteromedial thalamus; p, primary; s, secondary; d, dorsal; e, external; v, ventral.

LME ANOVA indicated a significant effect of state and a group x state interaction on node clustering coefficient across several ROI nodes. The nodes are summarized in **Figure 7C**. The nodes with group x state interactions included left anterior cingulate (F_24,380_=3.5, FDR p = 0.00001), left primary visual area (F_24,380_=2.6, FDR p = 0.003), left temporal cortex (F_24,380_=2.2, FDR p = 0.03), left bed nucleus of stria terminalis (BNST) (F_24,380_=2.3, FDR p = 0.01), right VPM thalamus (F_24,380_=3.2, FDR p = 0.00008). Nodes with main effect of state included the right pontine reticular formation (F_4,380_=6.1, FDR p = 0.003), inferior colliculus (F_4,380_=5.1, FDR p = 0.01), right SpVr (F_4,380_=6.4, FDR p = 0.003) and right central amygdala (F_4,380_=7.3, FDR p = 0.001).

LME ANOVA indicated a significant effect of state and a group x state interaction on node efficiency across several ROI nodes. The nodes are summarized in **Figure 7B**. The nodes with group x state interactions included left (F_24,380_=3.4, FDR p = 0.00002) and right anterior cingulate (F_24,380_=2.2, FDR p = 0.02), left insula (F_24,380_=2.0, FDR p = 0.04), left temporal cortex (F_24,380_=2.2, FDR p = 0.02), left primary visual area (F_24,380_=2.5, FDR p = 0.004), left BNST (F_24,380_=2.3, FDR p = 0.009), left vestibular nucleus area (F_24,380_=2.3, FDR p = 0.009), right piriform cortex (F_24,380_=2.0, FDR p = 0.04), right central amygdala (F_24,380_=1.9, FDR p = 0.04), right VPM thalamus (F_24,380_=3.6, FDR p = 0.000006). Nodes with main effect of state included the right central amygdala (F_4,380_=7.3, FDR p = 0.001), right pontine reticular formation (F_4,380_=3.3, FDR p = 0.00007), vestibular nucleus area (F_4,380_=4.9, FDR p = 0.02), right inferior colliculus (F_4,380_=4.3, FDR p = 0.04), right SpVr (F_4,380_=6.2, FDR p = 0.004) and right central amygdala (F_4,380_=8.5, FDR p = 0.0001).

### Aβ plaques distribute across cortical and subcortical areas that overlap with functional network states affected by amyloidosis

**Figure 7D-E** illustrates semi-quantitative whole brain distribution of Aβ plaque signal in 6-12 m TgCRND8 mice. Plaque burden is measured as percent occupation of ROI. Highest densities of plaques are localized to outer layers of the cortical mantle, with notably greater percent Aβ occupation in anterior cingulate, frontal cortical areas, somatosensory, motor, and auditory regions.

### Aβ plaque associated increases in DMN connectome strength and state fluctuations

We analyzed static and dynamic functional networks in an Alzheimer’s disease neuroimaging initiative (ADNI3) dataset (approved under University of Florida protocol #: NH00047739). Out of 34 scans, 27 were controls and 7 had subjective memory complaints. None were diagnosed with mild cognitive impairment or Alzheimer’s.

Group ICA revealed several well-established functional connectivity networks (**Figure 8A-J**). Medial and lateral visual cortical areas, auditory, sensorimotor, visuospatial systems, executive control, and dorsal visual stream were observed, as previously reported(Beckmann et al., 2005). Group averaged functional connectomes showing nodal strength organization in brains of low and high Aβ subjects are shown in **Figure 8K-L**. No significant differences in global network metrics were observed between low and high Aβ groups (**Figure 8M**). One-way repeated measures ME ANOVA indicated a group x ROI interaction within the DMN (F_7.8,251.3_ = 3.0, p = 0.003). Post hoc analysis of nodes within the DMN indicated significantly greater node strength in precuneus (p=0.0003, q=0.02), temporal area of the DMN (p=0.004, q=0.046) and temporoparietal areas 3 (p=0.0007, q=0.02), 5 (p=0.0009, q=0.02) and 7 (p=0.001, q=0.02) of high compared to low Aβ (**Figure 8N**).

**Figure 8.**
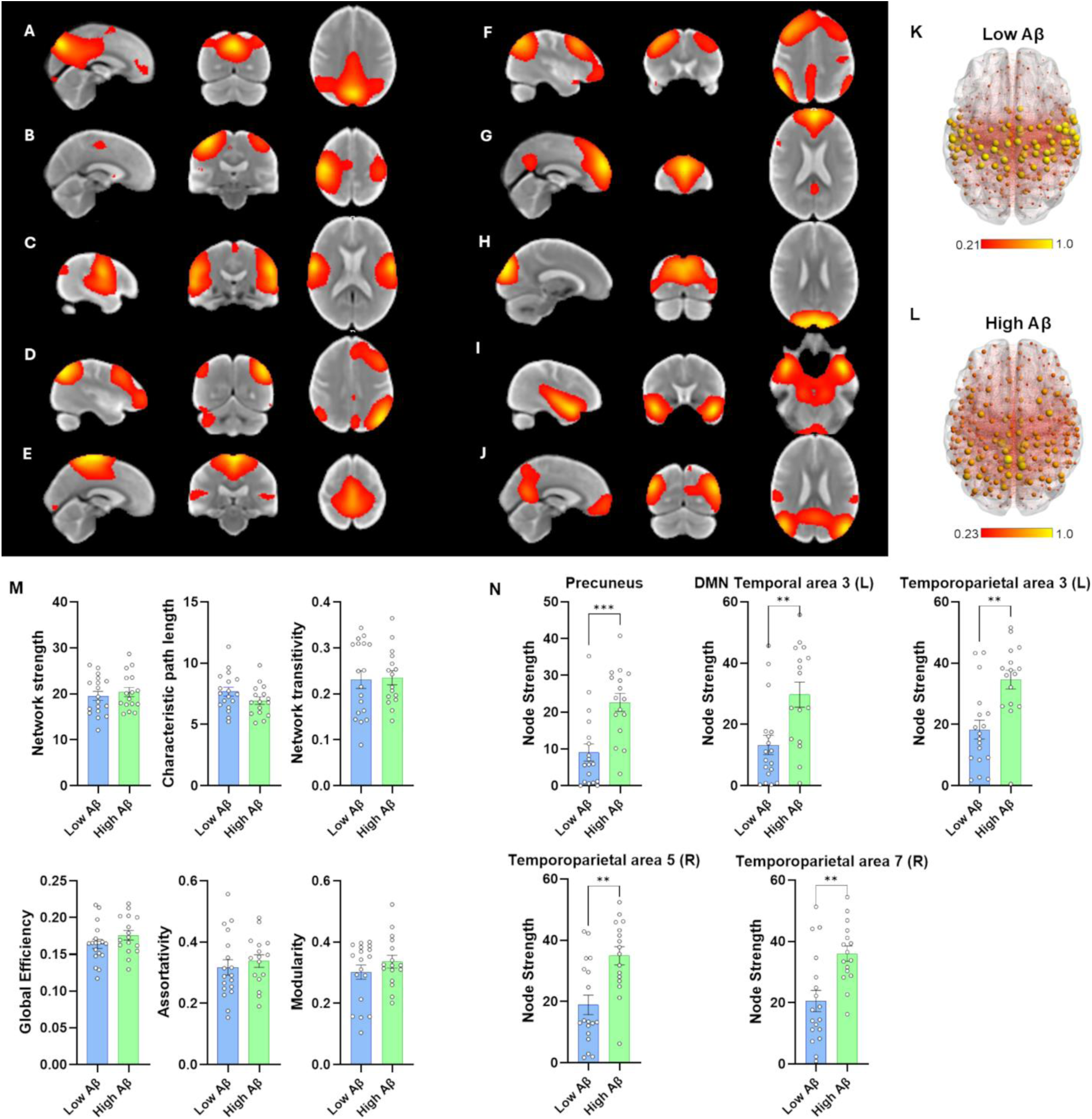
High Aβ plaque density is linked to greater node strength in default mode network areas. A) Default mode network areas. B) Right sensorimotor areas. C) Auditory areas. D) Left frontoparietal areas. E) Sensorimotor areas. F) Right frontoparietal areas. G) Executive areas. H) Medial visual areas. I) Auditory areas. J) Lateral visual areas. K) Group averaged functional connectome of low Aβ. L) Group averaged functional connectome of high Aβ. M) Global network measures. N) Connectivity strength in nodes of the default mode network. All data presented as mean ± standard error with overlaid scatter plots. Significant differences tested across all nodes using mixed effects ANOVA (FDR corrected). Post hoc Dunn’s tests indicated by asterisks (**p<0.01, ***p<0.005).

Mean FO across subjects was 0.33 ± 0.009 and mean dwell windows, 40.6 ± 1.1, supporting stable state durations. While maximum FO did not differ between groups, high Aβ subjects had greater switch rates between than low Aβ subjects (Mann-Whitney test p = 0.03) (**Figure 9A-B**). Thus, Aβ subjects had faster rates of switching between states. In contrast to the effect of Aβ on transition probabilities in mice, no differences in state transition probabilities were observed between low and high Aβ. **Figure 9C** shows mean functional connectome maps for low and high Aβ groups. As observed with static connectomes, a greater node strength is observed in high Aβ relative to low in posterior medial and lateral temporal and parietal cortical nodes within broader DMN areas. A group x state interaction was observed for FO, with greater state 1 FO in high vs low Aβ (2-way ME ANOVA; F_1.68,80.7_ = 5, p = 0.013; post hoc test p = 0.0028 (FDR adjusted), state 1 low vs high Aβ). A group x state interaction was observed for dwell windows, with lower values observed in high vs low Aβ (dwell windows: F_1.8,58.6_ = 6.7, p = 0.003; state 1 and 2) (**Figure 9D**).

**Figure 9.**
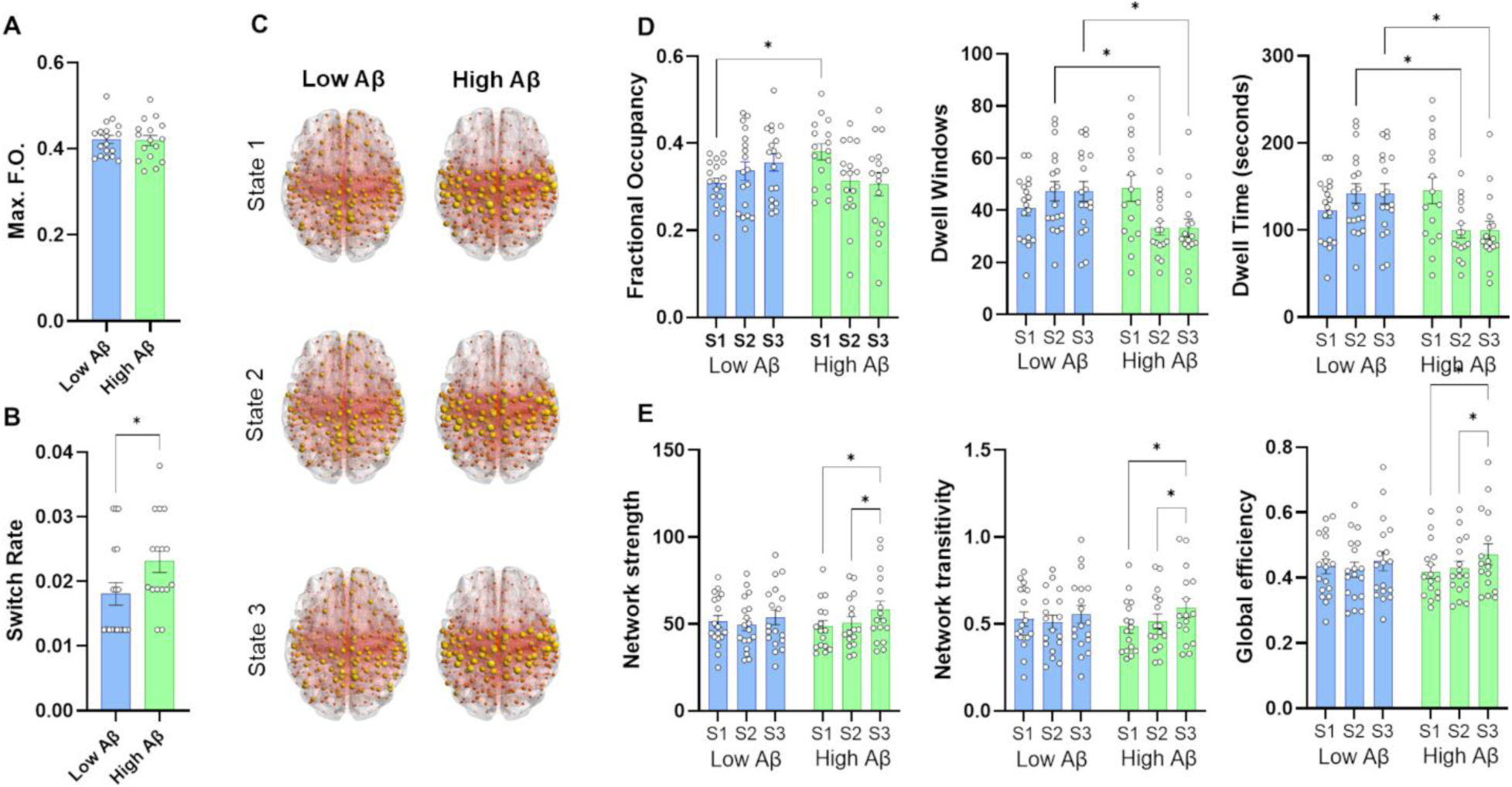
High amyloid burden is associated with high rates of state switching and instability of network strength, transitivity, and global efficiency across states. A) Maximum fractional occupancy per subject. B) State switch rates. C) Group-averaged dynamic functional connectome maps across 3 states for low and high Aβ. E) Network strength, transitivity, and global efficiency. All data presented as mean ± standard error with overlaid scatter plots. Significant differences tested across all nodes using mixed effects ANOVA (FDR corrected). Post hoc Dunn’s tests indicated by asterisks (*p<0.05).

A 2-way ME ANOVA indicated a main effect of state on node network strength, with high Aβ subjects having higher network strength in state 3 vs state 1 and 2 (F_1.8,60_=9.0, p=0.0005; Tukey’s post hoc test p = 0.01 state 3 vs 1, p = 0.03 state 3 vs 2). A similar effect of high Aβ was observed on network transitivity and global efficiency (transitivity: F_1.8,60_=9.2, p=0.0004; Tukey’s post hoc test p = 0.006 state 3 vs 1, p = 0.04 state 3 vs 2; efficiency: F_1.8,60_=10.2, p=0.0006; Tukey’s post hoc test p = 0.01 state 3 vs 1, p = 0.03 state 3 vs 2) (**Figure 9B**). LME ANOVA indicated node level differences between low and high Aβ that did not survive FDR correction.

## DISCUSSION

Our results indicate that Aβ is associated with increased static functional connectivity and network integration in both mice and humans but has dissociable effects on dynamic network organization across species. Specifically, dynamic network topology remained stable across states in Aβ mice, whereas human subjects with high Aβ exhibited increased state switching and altered state-dependent network measures. These results suggest that Aβ alone is insufficient to reproduce the dynamic network instability observed in human preclinical Alzheimer’s disease.

The present data agree with previous studies reporting increased functional connectivity between DMN-like areas in transgenic AβPP-bearing AD mice(Shah et al., 2016; Shah et al., 2018). We observed differences across TG and NTG groups, with the former having on average high connectivity strength, global efficiency and transitivity. At the nodal level, differences in these network measures were observed in anterior cingulate, retrosplenial cortex, subiculum, and primary visual areas, which are regions critical to cognitive and sensory processing, and that are known to accumulate Aβ plaques. Shah et al(Shah et al., 2016) were among the first to report hypersynchronous BOLD in both Tg2576 and PDAPP mice in areas that were also found here to have high node strength in middle aged TgCRND8 and 5XFAD mice. This was observed during pre-plaque stages in Tg2576 and PDAPP mice, both strains having single AD mutations. At later stages, functional connectivity declined significantly in their study(Shah et al., 2016), whereas in our study we observed elevated node strength in several regions in young TgCRND8 mice that already contain plaque deposits as early as 3mo and dense core plaques by 5mo (Chishti et al., 2001). Interestingly, node strength was mostly increased in middle age 5XFAD mice (15-19mo), which have elevated brain Aβ plaques and impaired contextual fear memory at 14mo(Neuner et al., 2019). The elevated connectivity strength persisted across more areas in both TG strains during 15-19mo. This is an important distinction between the studies since all strains produce human mutant Aβ and develop plaques, albeit at different ages and at different rates. TgCRND8 mice have double mutations while the diversity-variant 5XFAD we used here is on a mixed F1 B6D2 strain that harbors five AD mutations. An important distinction is thus that the PDAβPP and Tg2576 have single mutations, and in the case of the latter strain, these have vascular Aβ (as does the TgCRND8) and develop plaques more slowly. Still, this distinction does not explain why in our study we observed only elevated node strength, even at the later age group of TG mice.

The discrepancy may be due to the analysis approach, with Shah et al(Shah et al., 2016) analyzing single pairwise functional connectivity which differs from our calculation of node strength. The approach used by Shah et al(Shah et al., 2016) is a specific readout for pairwise changes between two nodes over time, whereas node strength can remain elevated for one node if its pairwise connectivity is strong with any other set of nodes in a network. One is physiologically grounded and the other based on a statistical measure of node ‘prominence’ within a network (strength of overall ‘ties’ within a network). Work by Grandjean and colleagues(Grandjean et al., 2016) evaluated the effects of vascular, parenchymal, and intracellular Aβ in 3 strains of amyloid mice that preferentially deposit plaques in these compartments(Grandjean et al., 2016). Within and between ICA component analysis was carried out, which showed reduced within-component functional connectivity (including cingulate-ventral hippocampus) and increased between-component connectivity (including prefrontal-cingulate). This finding supports the notion of reorganization of functional connectivity across groups of nodes, perhaps as part of the compensatory response to Aβ accumulation. It also should be noted that additional work by Shah et al(Shah et al., 2018) correlated increased pre-plaque functional connectivity with spatial learning deficits, based on performance on a Morris Water Maze task in the AβPP^NL-F^ knock-in mouse (Swedish/Iberian double mutations)(Shah et al., 2018). A limitation of the present work is that we did not include behavioral assays for cognitive performance to evaluate connectome measures as a function of cognition behaviors.

To our knowledge, there have been no studies to date that have examined how Aβ affects HMM-derived fMRI connectivity network states, a question that can be feasibly studied in mice engineered to express Aβ. We failed to observe significant effects of Aβ on network measures across HMM states, and the latter appeared relatively stable in terms of time spent in each state and switching between states. On the other hand, analysis of quasi-periodic patterns (QPPs), known to contribute to spatial and temporal dynamics in resting state fMRI(Thompson et al., 2014), have been shown to be affected by Aβ and age. This points to potentially distinct mechanisms driving QPPs versus the observed HMM states. Middle aged (18mo) Tg2576 mice that develop Aβ plaques around 9-11 mo had lower occurrence rates for QPPs compared to age-matched controls (Belloy et al., 2018). DMN-like areas also showed reduced anticorrelated functional connectivity with task-positive areas of the mouse brain (Belloy et al., 2018). Our results suggest that the effect of Aβ on static network measures is dissociable from the dynamic functional network stability that was observed in mice. The relatively stable state patterns may be due to lack of behavior or the effects of sedation, which is not the case of human functional connectomes. However, unknown pathologies present in human connectomes, and not in our Aβ mice, may also explain our results. Combining AD-relevant pathologies with other known non-pathological factors (e.g., diet, exercise, social interactions, chronic stress or trauma) may bring mouse dynamic connectomes closer to representing the state dependent variations observed in human subjects.

Human functional connectome results are consistent with previous reports. In terms of static network strength, we observed increased node strength in precuneus, and other regions of the DMN. Previous research indicated that Aβ (and tau) in cognitively unimpaired subjects (∼60 years of age) is associated with increased functional connectivity between precuneus and anterior hippocampus(Fischer et al., 2025). This result is consistent with in the observed increases in strength, efficiency and transitivity in DMN-like areas of Aβ mouse static networks. As in mouse networks, the effect of Aβ plaques may include neuronal, synaptic, and vascular deficits that can explain the present results. Pathological changes in response to Aβ could provide the basis for within-state changes in network measures, where some states are visited more often than others. A reduction in frequency of visits to one of three identified states was observe in AD (females but not males)(Sendi et al., 2023a). This differs from our results, which included mostly women, and were pre-selected for not having signs of cognitive impairment. Visits to specific states vary as a function of dementia risk factors and self-reports of cognitive performance(Dautricourt et al., 2022). Interestingly, frequent visits to a single state are observed in small vessel disease(Chen et al., 2024; Mao et al., 2025) and appear to distinguish small vessel disease from AD(Fu et al., 2019). This suggests a role for either changes in small vessel structure and function or a role of astrocytes that regulate neurovascular coupling and local perfusion.

Previous work using K means clustering determined that fMRI timeseries in mice could be segmented into 6 states (co-activation patterns) with similar network features(Gutierrez-Barragan et al., 2022). Such an approach captures rodent network features that may be conserved and translatable to primate networks(Gutierrez-Barragan et al., 2024). Twenty distinct explanatory vectors were identified using a dictionary learning algorithm, which also identified relationships of several of these states (atoms) to chronic stress exposure(Grandjean et al., 2017). This approach in mice may model aspects of human studies identifying links between dynamic functional connectivity and neuropsychiatry condition-specific network states(Rashid et al., 2014). The HMM variant used here applied a mixture of Gaussians under variational Bayes framework to find a lower evidence bound solution to segmenting network-based time series into predefined states across all subjects. Thus, while it was found that the networks could be segmented into 5 states but not 6 or greater, this was determined across the entire cohort of mice that varied by age and Aβ conditions (and by sex). Thus, a more uniform cohort of mice may may produce a different outcome (how many states are segmented). Further, the inclusion of behavioral assessments(Benisty et al., 2024), especially close enough in time to the imaging as in previous work in rats(Pompilus et al., 2020), may raise the predictive validity of these approaches to distinct behavioral states(Shahsavarani et al., 2023).

There are several limitations in our study that should be noted. First, the use of anesthetic and sedative are sure to impact what aspects of dynamic network connectivity are captured in mice(Kang et al., 2018; Tsurugizawa and Yoshimaru, 2021; Gutierrez-Barragan et al., 2022). The protocol used here, however, is often used across laboratories and offers a good solution to image sedated mice and comparing results across mouse imaging laboratories. Awake mouse studies are feasible and emerging protocols for training mice during MR scanning may facilitate interpretations when analyzing dynamic functional connectivity networks(Bergmann et al., 2020; Gutierrez-Barragan et al., 2022; Brems et al., 2026). Recent work using a simple motor task along with the use of the movement-insensitive zero TE sequence seems like a strong future direction(Daley et al., 2026). Implementation of this approach, or similar methods, across preclinical laboratories should improve harmonization of data collected in awake mice. Future studies should include a variety of phenotypes and pathologies to ensure statistical robustness in capturing the effects Aβ relative to other factors.

## Acknowledgements

This work was funded by NIA R21AG065819 with additional support from the McKnight Brain Institute of the University of Florida and a AI2020 Catalyst Grant (AWD09459). Neuroimaging of mice was performed in the National High Magnetic Field Laboratory’s AMRIS Facility, which is funded by National Science Foundation Cooperative Agreement No. DMR-1644779 and the State of Florida.

## Supporting Information Figures

**Supplemental Figure 1.**
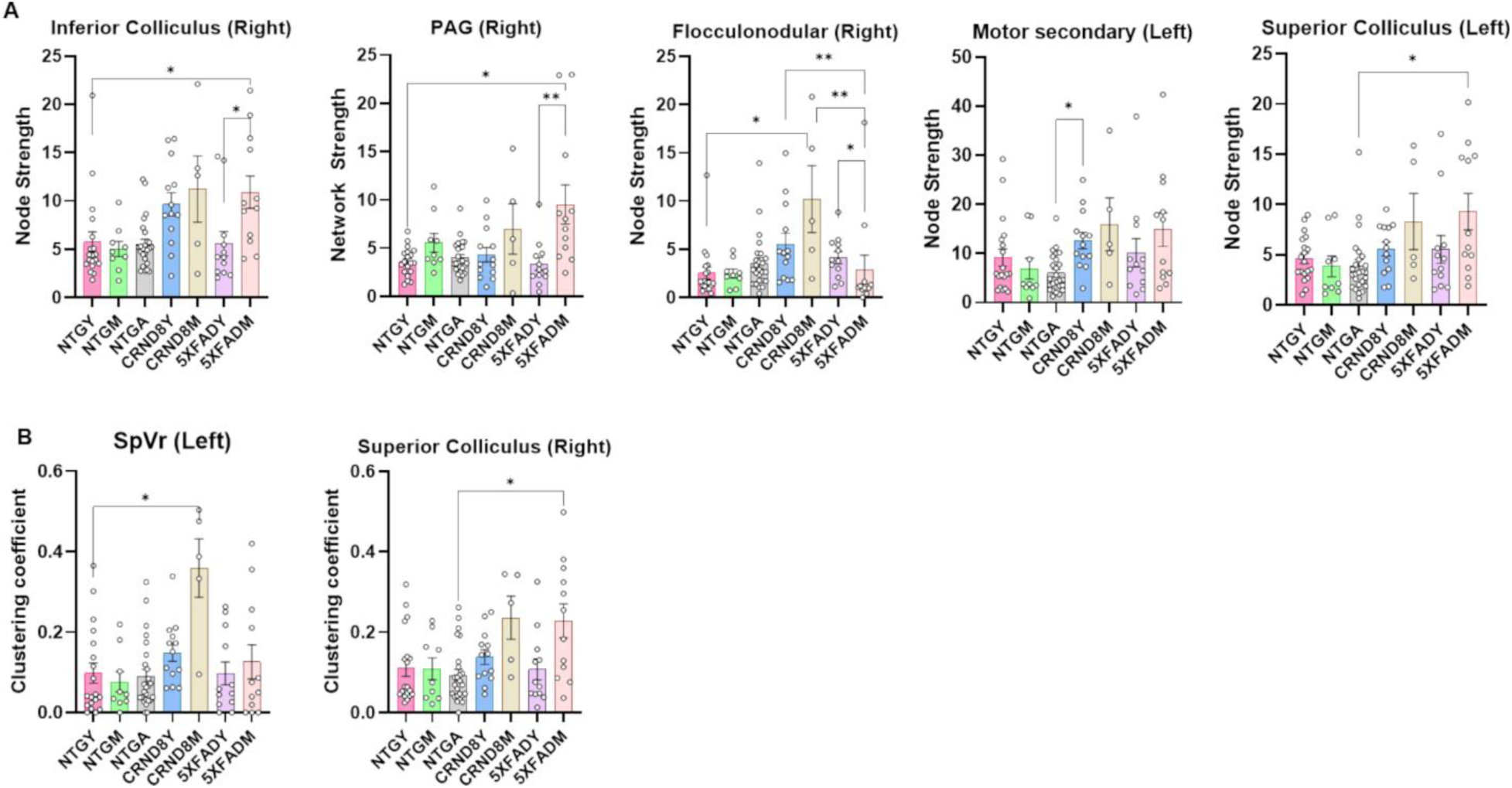
Node strength and clustering coefficient in additional brain areas affected by amyloidosis. Groups are as shown in Figure 2. A) Node strength. B) Clustering coefficient. All data presented as mean ± standard error with overlaid scatter plots. Significant differences tested across all nodes using linear mixed effects ANOVA (FDR corrected). Post hoc Dunn’s tests indicated by asterisks (*p<0.05, **p<0.01).

